## Supplemental figures S1-S7 and supplemental movie legends for "Cell morphology and nucleoid dynamics in dividing *D. radiodurans*"

[1] Univ. Grenoble Alpes, CEA, CNRS, IBS, F-38000 Grenoble, France.

[2] Institute for Integrative Biology of the Cell (I2BC), CEA, CNRS, Univ. Paris-Sud, Université Paris-Saclay, Gif-sur-Yvette, France.

[3] Univ. Grenoble Alpes, CNRS, DPM, 38000 Grenoble, France.

[4] These authors contributed equally.

[*] Correspondance:

**Supplemental Information**

Supplemental Figures S1-S7

Supplemental Movies S1-S5

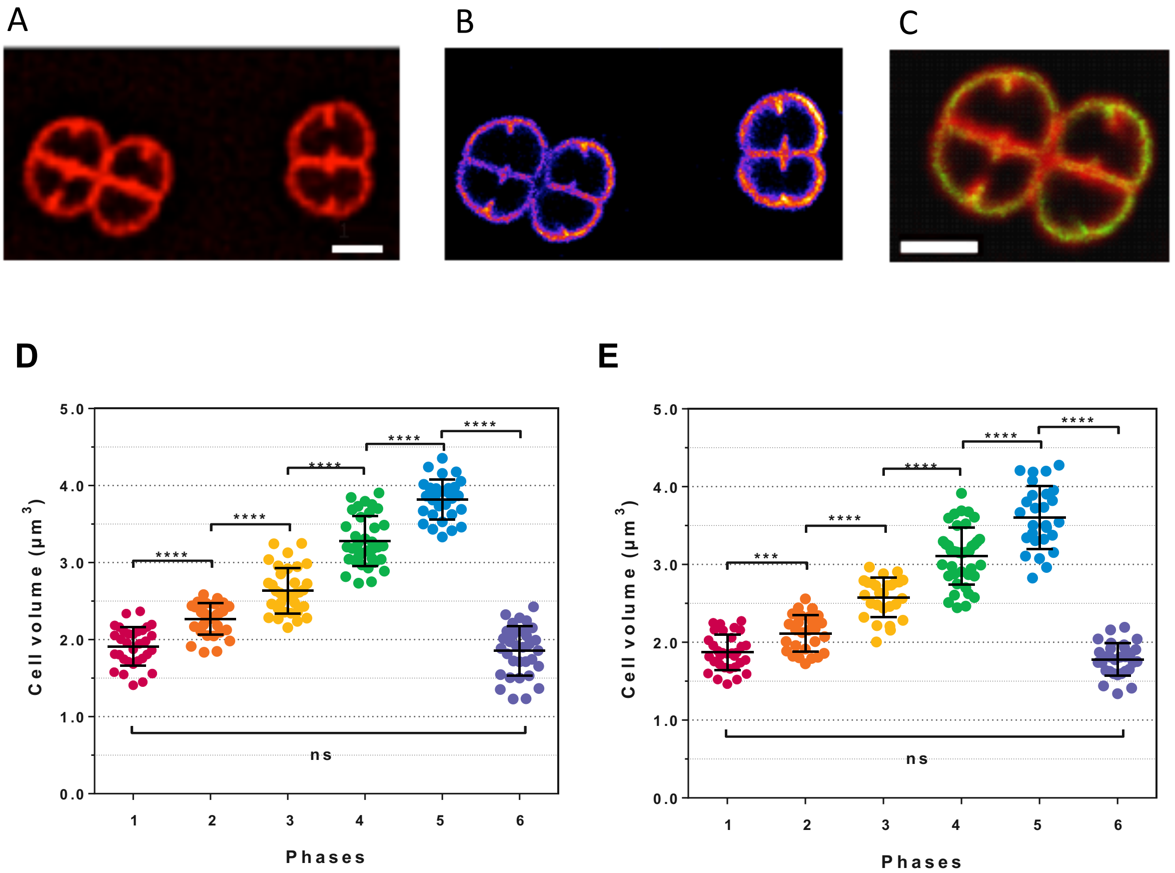


**Figure S1: Comparison of spinning-disk and PAINT images of *D. radiodurans* cells stained with the membrane dye Nile Red.** (A) Spinning-disk image and (B) PAINT image of the same Nile Red stained *D. radiodurans* cells. (C) Overlay of images (A) and (B). Scale bar: 1 µm. (D)-(E) Changes in cell volume as a function of the cell cycle retrieved from either super-resolved, PAINT images (D), or from spinning-disk confocal images (E) of Nile Red stained *D. radiodurans*. The same three independent populations of cells were observed with the two imaging techniques. (N=180 cells, N>27 for each phase). Data are represented as mean ± SD. Individual values are shown as dots. Cell volumes were calculated by measuring the cell parameters as described in Figure S2. Only very small differences in cell volumes were observed between the two imaging methods. ***: P<0.001, ****: P<0.0001, ns: not significant.

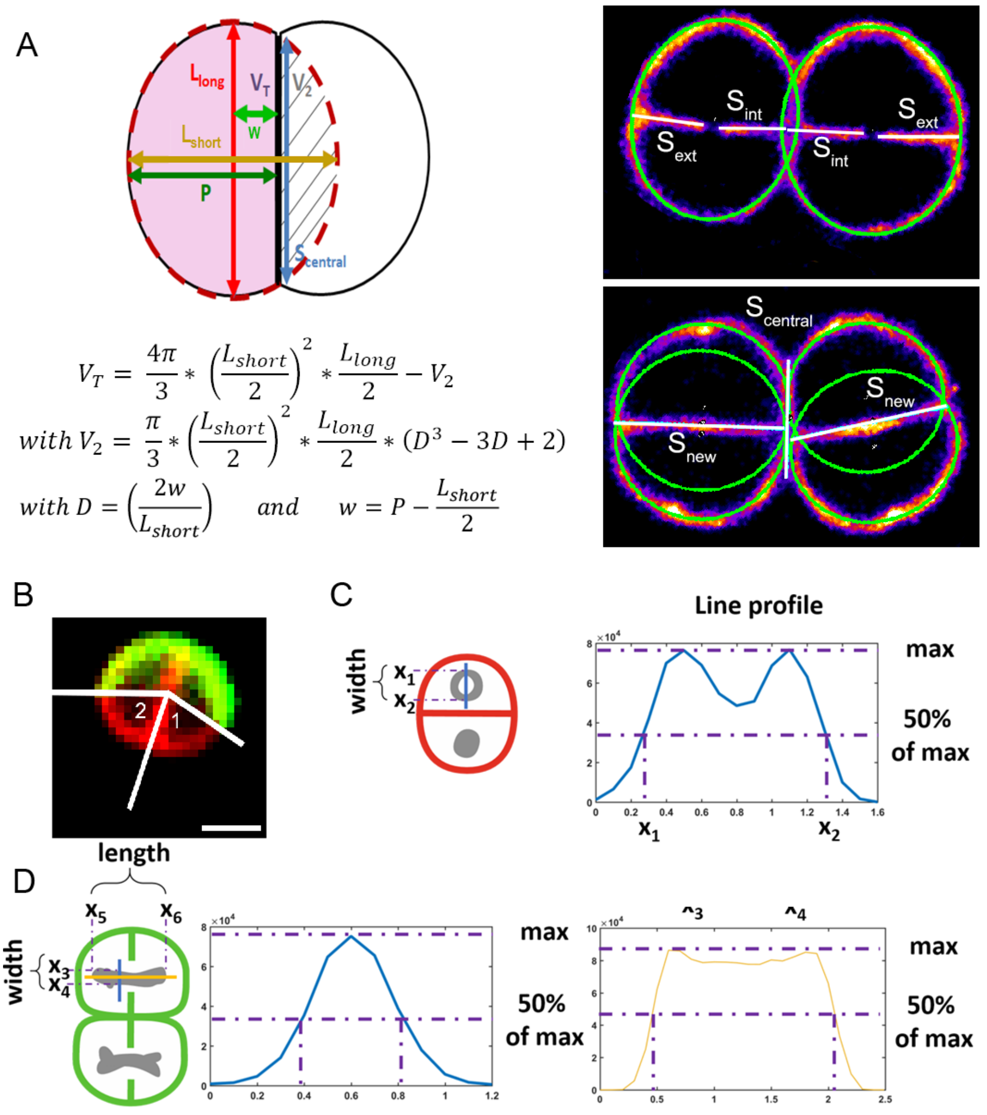


**Figure S2: Cell and nucleoid measurements used in this study.** (A) Schematic diagram illustrating the mode of calculation of the volumes of *D. radiodurans* cells based on the fitting of ellipses (dotted red line) to individual cells and the equations used for these calculations. $L_{long}$= length of major axis of the fitted ellipse; $L_{short}$= length of minor axis of the fitted ellipse; P= distance between central septum and the opposite side of the ellipse; w= distance from center of ellipse to central septum (=P-[$L_{short}$/2]); $S_{ext}$=length of exterior septum of cells in Phases 4 and 5; $S_{int}$= length of interior septum of cells in Phases 4 and 5; $S_{new}$= length of the newly formed central septum in Phase 6; $S_{central}$ = length of the old division septum originating from the previous cell cycle. (B) Illustration of the two angle measurements made to determine the lengths of the BADA (green) and Nile Red (red) labelled cell perimeters in Phase 1 diads at T= 130 min. Scale bar: 1 µm. (C)-(D) Illustration of the measurements made to determine the width of the toroidal-shaped nucleoids (C) and the length and width of the elongated nucleoids (D).

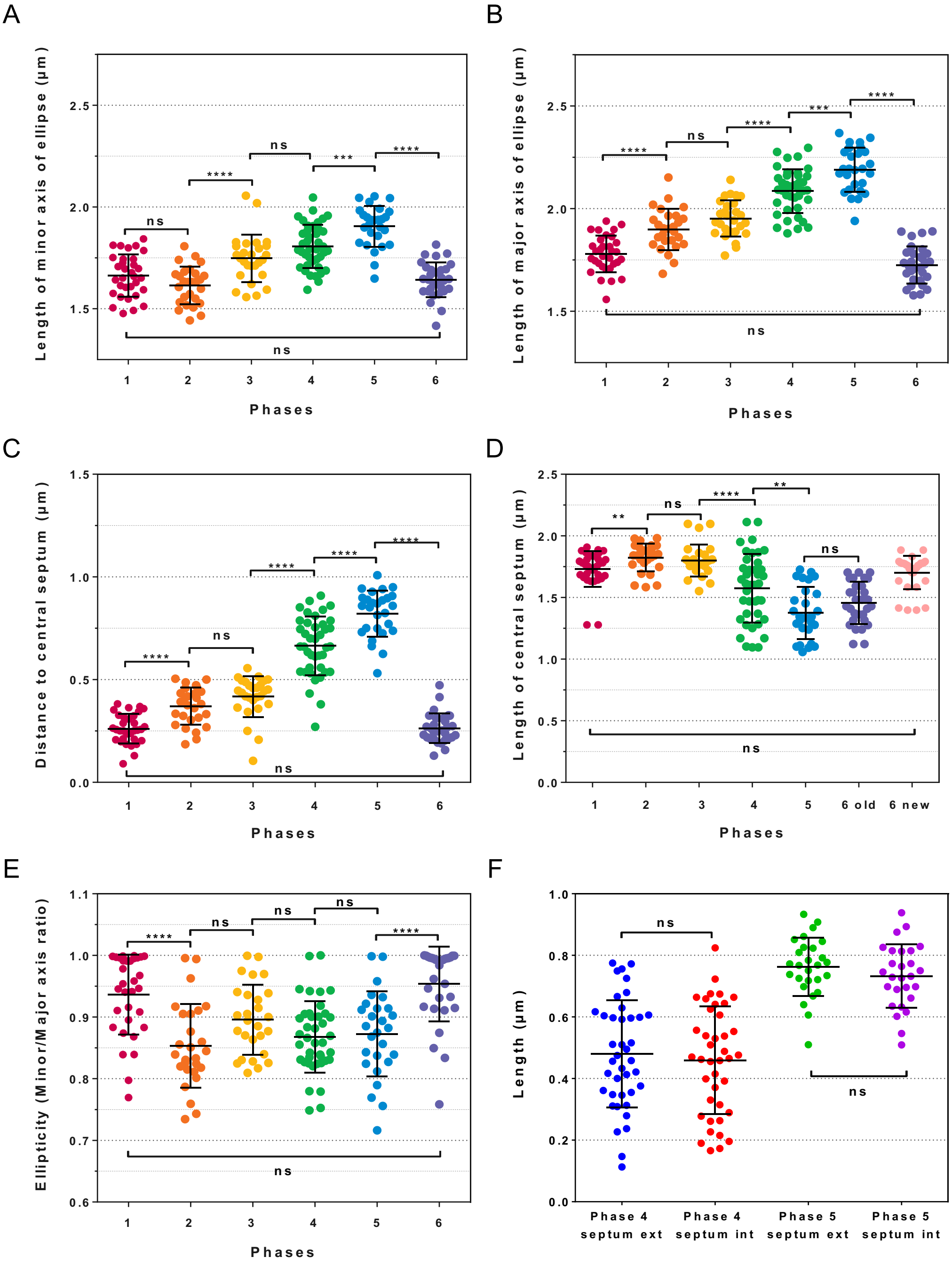


**Figure S3: Cell parameters of *D. radiodurans* cells extracted from PAINT images of Nile Red stained, exponentially growing cells.** (A)-(B) Length of the minor (A) and major (B) axes of the fitted ellipses used to measure cell volumes (see Figure S2). (C) Distance of the ellipse centre to the central septum of the cell. (D) Length of the central septum. In Phase 6 cells, there are two central septa: the old septum originating from the previous cell cycle and the new one that has just closed. (E) Ellipticity of the cells (ratio between the length of the minor and major axes). (F) Lengths of the outer (growing from the peripheral cell wall) and inner (growing from the central septum) septa in Phases 4 and 5 (see Figure S2). N=180 cells, N>27 for each phase. Data are represented as mean ± SD. Individual values are shown as dots. *: P<0.05, **: P<0.01, ***: P<0.001, ****: P<0.0001, ns: not significant.

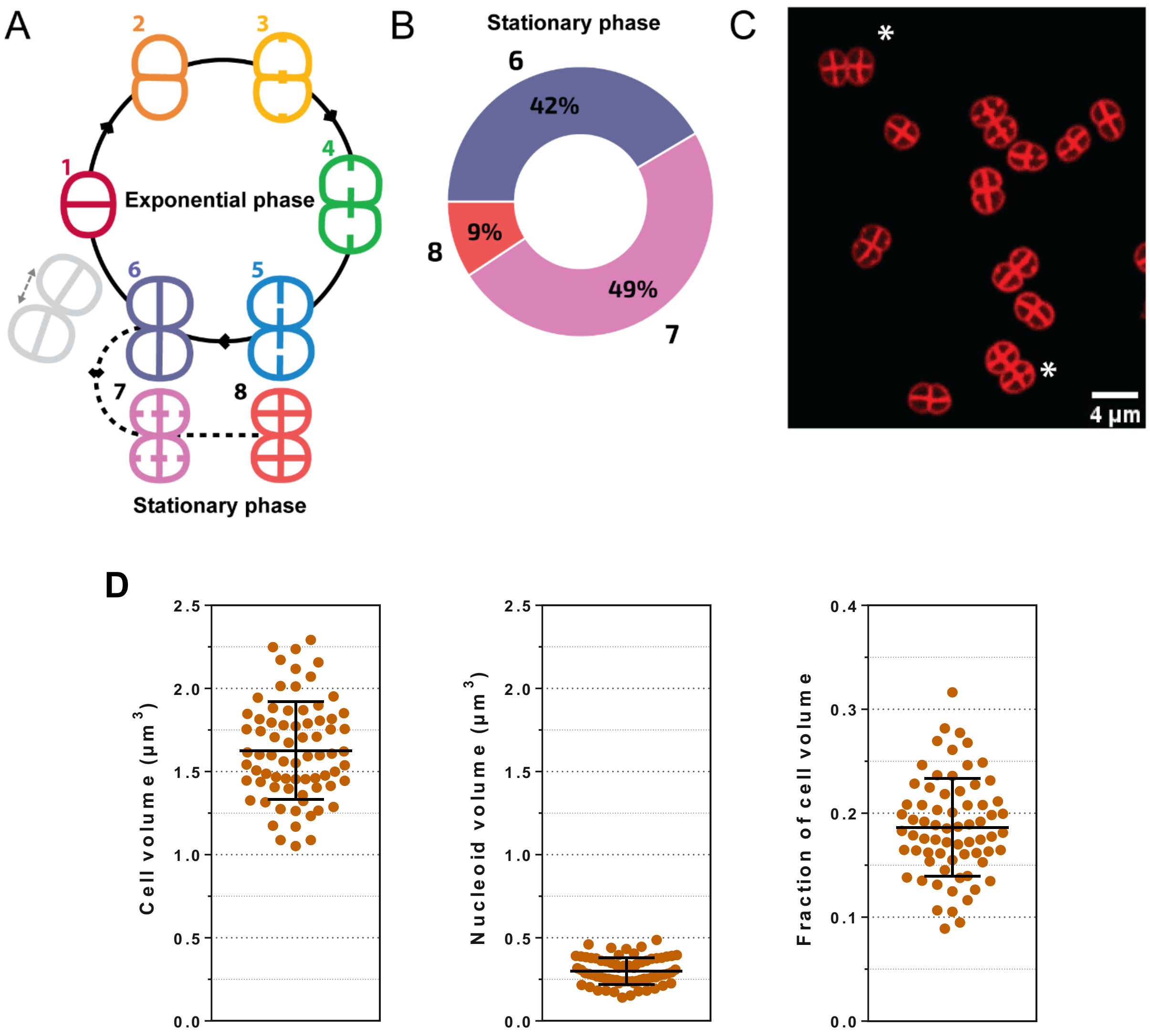


**Figure S4: Changes in cell morphology during the stationary phase of *D. radiodurans* cell growth**. (A) Schematic representation of the cell cycle, for one diad. In stationary phase, two additional phases can be seen: Phase 7 corresponds to tetrads engaging in a new cell cycle without dissociation into diads, and Phase 8 cells are octads. (B) Distribution of phases in the population of stationary cells (24h of growth) when observed at a given time point (N>600). All data were collected from at least 2 independent experiments. (C) Spinning-disk image of Nile Red stained *D. radiodurans* stationary cells. Octad cells are marked with an asterisk. Scale bar: 4 µm (D) Mean cell and nucleoid volumes, and fraction of the cell volume occupied by the nucleoid in stationary phase 6 tetrads (N=70 cells). Cell volumes were calculated by measuring the cell parameters presented in Figure S2 on spinning-disk images. Nucleoid volumes were measured as described for exponentially growing *D. radiodurans* cells (see Methods). Data are represented as mean ± SD. Individual values are shown as dots.

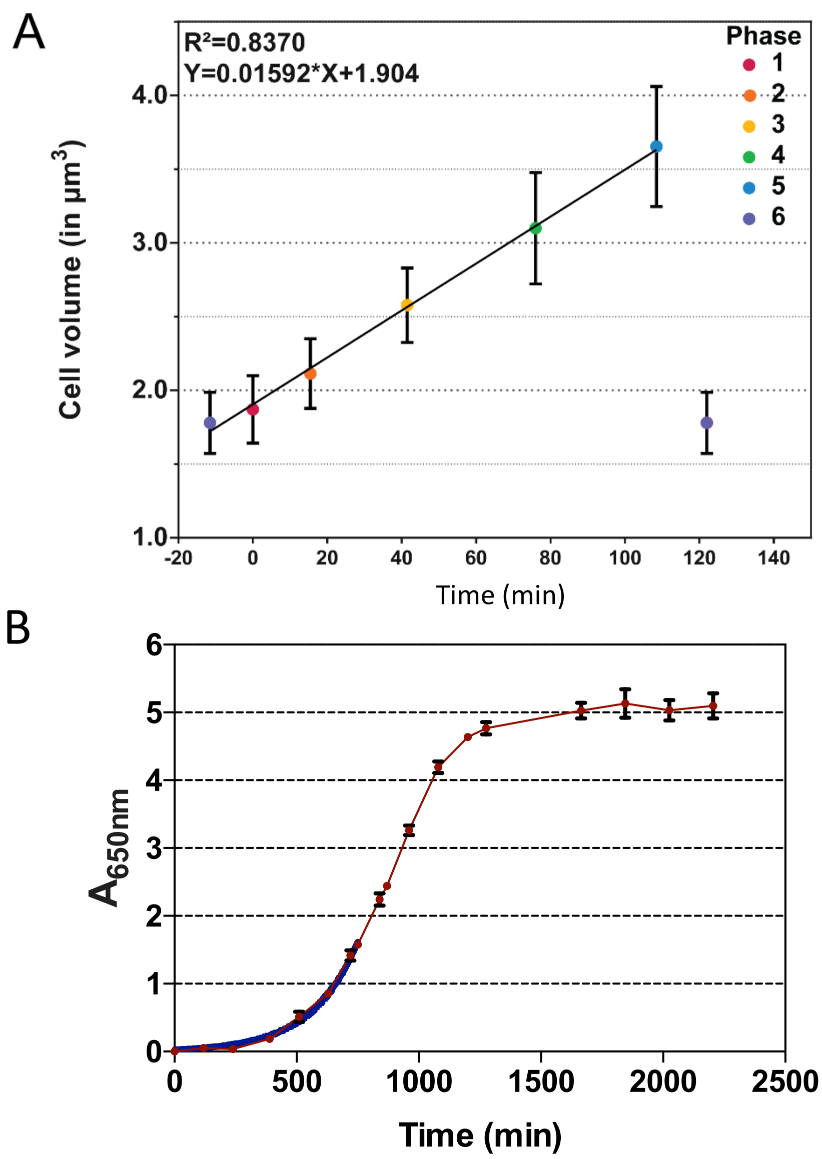


**Figure S5 - Related to Figure 1: Changes in cell size and optical density as a function of incubation time.** (A) Linear increase of *D. radiodurans* cell volume with time during the cell cycle, derived from measurements of Nile Red stained exponentially growing cells imaged with PAINT. (B) Growth curve of wild-type *D. radiodurans* cells grown in TGY2X medium at 30°C in a shaking incubator. The curve was established using data points from at least three independent experiments. The doubling time of exponentially growing cells was derived by fitting the initial growth phase to an exponential growth curve (blue; see Methods) and was estimated to be 130 minutes.

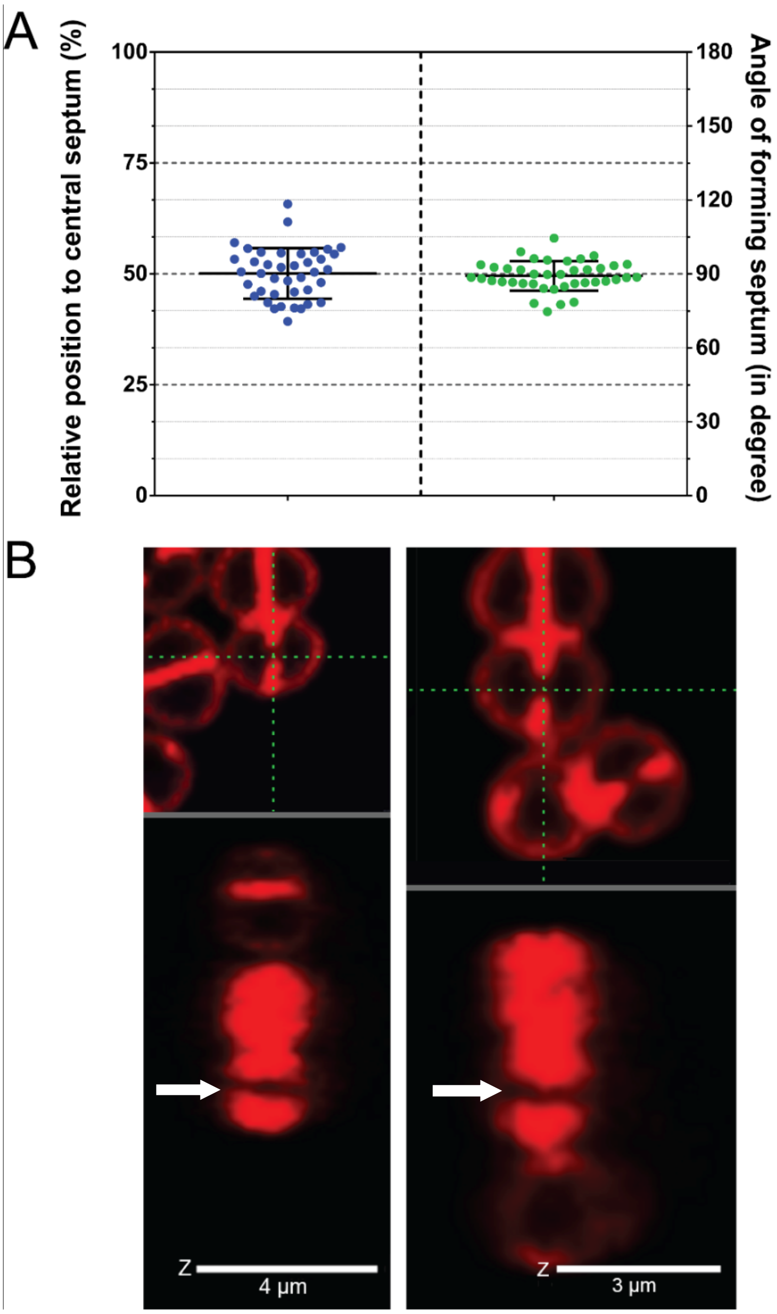


**Figure S6 - Related to Figure 2: Septal growth and closure mechanism in *D. radiodurans*.** (A) Position and angle of newly forming septa relative to the central septum (N>40 cells)**.** Data are represented as mean ± SD. Individual values are shown as dots. (B) Two examples (left and right) of 3D spinning-disk images of Nile Red stained *D. radiodurans* cells in which septal closure is in progress. The top panel represents the observed cells, in the XY plane, with the observation cross superimposed in green. The bottom panel is the XZ orthogonal view along the X direction defined by the green cross in the top panel. Septal growth from both sides of the cells leaves a gap stretching all away across the height of the cell, as seen in the XZ projection and indicated with white arrows. These images support a closing door mechanism rather than a diaphragm.

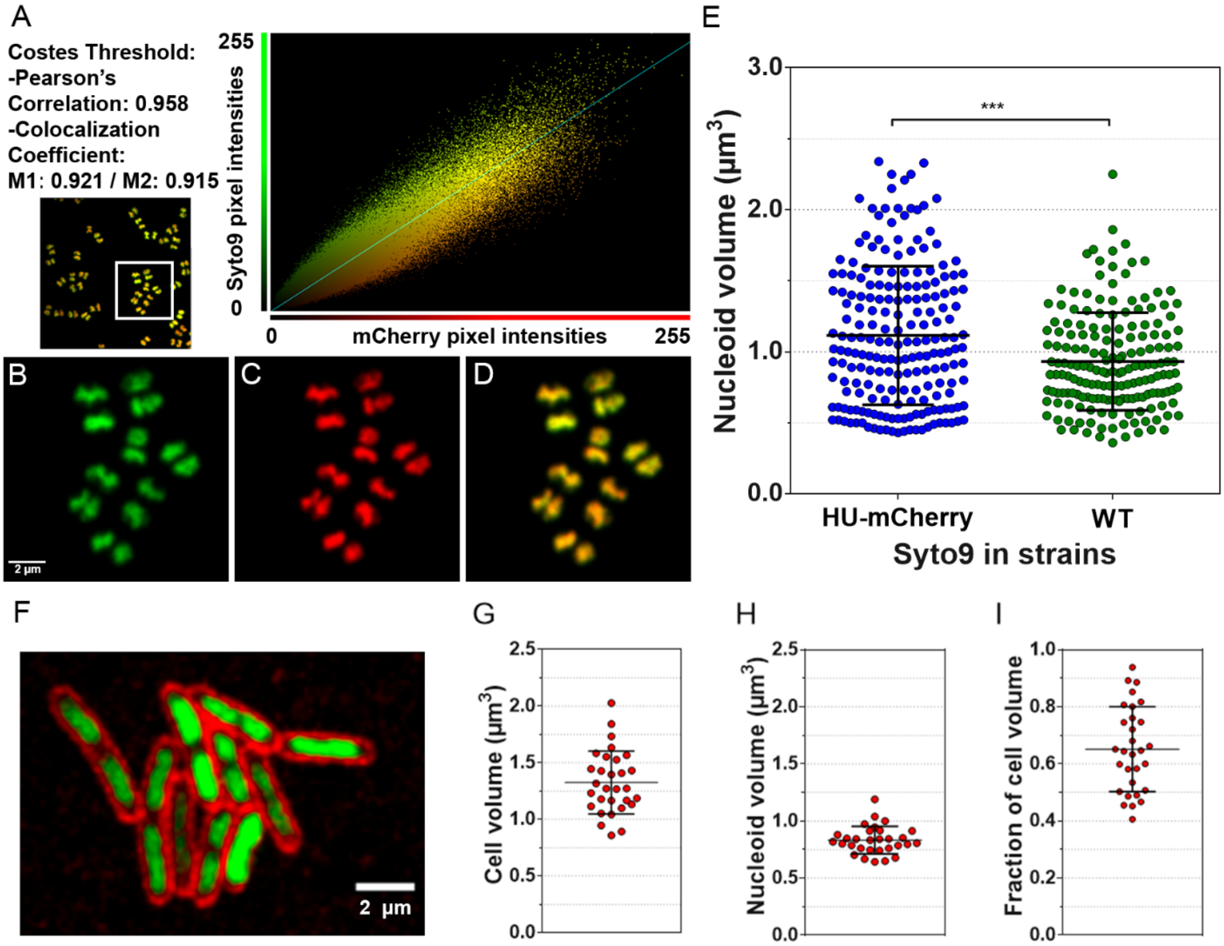


**Figure S7 - Related to Figures 3 and 4: Nucleoid labelling in exponentially growing *D. radiodurans* and *E. coli* BL21 cells.** (A) Colocalization statistics (analysis with Volocity software) of HU-mCherry and Syto9 fluorescence signals in *D. radiodurans* cells expressing HU-mCherry and stained with Syto9. The colocalization analysis was performed on the full Z-stack of untreated raw images. (B)-(D) Close-up insets of white box in (A), showing nucleoids stained with both Syto9 (B) and HU-mCherry (C). (D) Overlay of the two fluorescence signals. Similar shapes and structures are seen using both labelling methods. The images correspond to raw images, shown in extended focus (all Z-planes are superimposed into a single, final image). (E) Nucleoid volumes of Syto9 stained exponentially growing wild-type (WT) and HU-mCherry expressing *D. radiodurans* cells (N>200; ***: P<0.001). Data are represented as mean ± SD. Individual values are shown as dots. (F) Image of exponentially growing *E. coli* BL21 cells stained with Syto9 and Nile Red. (G) Cell volume, (H) nucleoid volume and (I) fraction of the cell volume occupied by the nucleoid (N>30) in *E. coli* cells. The fraction consists of the ratio of the nucleoid volume divided by the volume of the associated cell. The cell volume was computed from measurements of the length and width of individual cells, assuming *E. coli* cells were cylinders capped with two semi-spheres. The nucleoid volumes were extracted as for *D. radiodurans* nucleoids (see Methods). Data are represented as mean ± SD. Individual values are shown as dots.

**Movie S1: Growth and division of *D. radiodurans.***

Time-lapse movie of live, Nile Red stained, exponentially growing *D. radiodurans* cells, deposited on an TGY2X agarose pad. Images were acquired every 10 min for a total period of 4 h. Scale bar: 2 µm.

**Movie S2: Nucleoid dynamics in live, wild-type *D. radiodurans*.**

Time-lapse movie of live, Syto9 stained, *D. radiodurans* cells, deposited on an TGY2X agarose pad. Images were acquired every 10 min for a total period of 3 h. Scale bar: 5 µm.

**Movie S3: Nucleoid dynamics in live, HU-mCherry expressing *D. radiodurans*.**

Time-lapse movie of live, HU-mCherry expressing *D. radiodurans* cells, deposited on an TGY2X agarose pad. Images were acquired every 10 min for a total period of 3 h. Scale bar: 5 µm.

**Movie S4: Simulation of the major morphological changes occurring at the cellular and nucleoid level in dividing *D. radiodurans.***

On-scale (size and timing) simulation of *D. radiodurans* cell cycle, illustrating the coordination of the morphological changes of the nucleoids with septal growth during the cell cycle. This movie was created based on the cell and nucleoid measurements presented in Figures 1D, Figure 4A and Figure S3 and on the cell cycle duration presented in Figure 1E. Scale bar: 1 µm.

**Movie S5: Minute-scale dynamics of *D. radiodurans* nucleoids.**

Time-lapse movie of live, HU-mCherry expressing *D. radiodurans* cells, deposited on an TGY2X agarose pad. Images were acquired every 20 sec for a total period of 10 min. Scale bar: 4.2 µm.
